# Palmitoylation is critically required for cancer intrinsic PD-1 expression and functions

**DOI:** 10.1101/625483

**Authors:** Han Yao, Chushu Li, Jean-Philippe Brosseau, Huanbin Wang, Haojie Lu, Caiyun Fang, Hubing Shi, Jiang Lan, Jie Xu

## Abstract

Programmed cell death protein 1 (PD-1) is a crucial anticancer target, but the relatively low response rate and acquired resistance to existing antibody drugs highlight an urgent need to develop alternative targeting strategies. Here we report the palmitoylation of PD-1, discovered the main DHHC enzyme for this modification, revealed the mechanism for its effect on PD-1 expression, and rationally developed a peptide for targeting PD-1 expression. Palmitoylation promoted the trafficking of PD-1 to recycling endosome, thus preventing its lysosome-dependent degradation. Palmitoylation was required for the activation of PD-1 downstream signaling, and targeting palmitoylation by pharmacological inhibitor or depleting the modification enzyme caused significant anti-tumor effects. A peptide was designed to competitively inhibit PD-1 palmitoylation and expression, opening a new route for developing PD-1 inhibitors as a strategy for cancer immunotherapy.

**Significance:** We show for the first time that PD-1 is palmitoylated, identify DHHC9 as the predominant enzyme for its palmitoylation, and reveal the molecular mechanisms underlying its effects on PD-1 stability and functions. Importantly, we also designed a competitive inhibitor targeting PD-1 palmitoylation, and this first-in-class molecule may inspire the development of new checkpoint inhibitors.

## Introduction

Programmed death protein 1 (PD-1) and its two natural ligands PD-L1 and PD-L2 deliver inhibitory signals to regulate the balance between T cell activation, tolerance, and immunopathology. PD-1 is expressed on the surface of activated T cells as an inhibitory receptor ^1^, while its ligands PD-L1 and PD-L2 are mainly expressed in antigen-presenting cells and tumor cells ^2^. PD-1 downstream signaling includes the suppression of T cell proliferation, cytokine production, and cytotoxic functions. Therapeutic antibodies targeting PD-1 have displayed remarkable anti-tumor efficacy, However, most patients do not show durable remission, and some tumors have been completely refractory to response with checkpoint blockade ^3^, highlighting a need for further understanding the regulation of PD-1 ^4^.

Recent studies revealed intrinsic expression of PD-1 in melanoma ^5^, liver cancer ^6^ and other cancers ^7^. In these cases, PD-1 can modulate mTOR signaling and promote tumor growth independently of the adaptive immune system ^5,6^. In tumor cells, PD-1 was found to promote tumor growth even in the absence of functional adaptive immune system, which involved the increased phosphorylation of ribosomal protein S6 (RPS6) and eIF4E as effectors of mammalian target of rapamycin (mTOR) signaling^5,6^.

Palmitoylation is typically the covalent attachment of palmitic acid to cysteine of membrane proteins, which serves as a mechanism to regulate protein localization and function^8^. Palmitoylation may be catalyzed by a family of aspartate-histidine-histidine-cysteine (DHHC) acyltransferases, which display different specificities to existing substrates such as Ras, EGFR, Wnt, and Shh^9^. Recently, we found that PD-L1 palmitoylation by DHHC3 stabilizes PD-L1 through the suppression of ubiquitination and lysosomal degradation ^10^. However, it has not been reported previously whether palmitoylation may play a role in the regulation of PD-1. Here we report that PD-1 is palmitoylated at Cys192 residue by the DHHC9 acetyltransferase, and this chemical modification promotes PD-1 expression and intrinsic signaling to promote cancer cell growth. These findings may bring new insights into the regulation of PD-1, and provide additional opportunity for drugging PD-1 as an important anti-cancer target.

## Results

### PD-1 is palmitoylated at Cys192

Palmitoylation is an established post-transcriptional modification regulating the abundance of various cancer-associated protein ^11^ but PD-1, a key immunocancer target, was never reported to be palmitoylated. To identify a putative S-palmitoylation site on PD-1, we performed an silico motif-based prediction using the MDDPalm algorithm ^12^. The result indicated that the Cys192 residue that joins the transmembrane domain and cytosolic part of PD-1, resembling a typical membrane proteins palmitoylation site and matched to a previously characterized palmitoylation motif ^13^ (**Fig.1a**). To directly experimentally validate the palmitoylation of PD-1, we performed a palmitoylation-specific pulldown assay by Click-iT labeling ^14^ on endogenous PD-1. Briefly, cultured cells were fed with azidopalmitate as a source of palmitic acid. Proteins were harvested and labeled with biotinalkyne, followed by streptavidin pull-down and immune-blot using an antibody specific to the protein of interest. (schematics in **Fig.1b**). We were able to successfully obtain a reliable signal when we used anti-PD-1 antibody in protein extracted from NB4 cells (**Fig. 1c**, upper panel) and Molt-4 cells (**Fig. 1d**). Similar results were obtained using an independent anti-PD1 antibody (**Fig. 1c**, lower panel). Thus, this assay confirmed the palmitoylation of endogenous PD-1 protein. To directly validate the palmitoylation of PD-1 at the residue Cys192, we engineered a mutant version of PD-1 at position 192 (Cys192Ser) and challenged its capacity to be palmitoylated by Click-iT chemistry. As expected, replacing the Cys192 for a serine blocked the palmitoylation of PD-1 (**Fig.1e**), indicating that the residue Cys192 is a palmitoylation site and most probably the only site for PD-1 palmitoylation.

**Figure 1.**
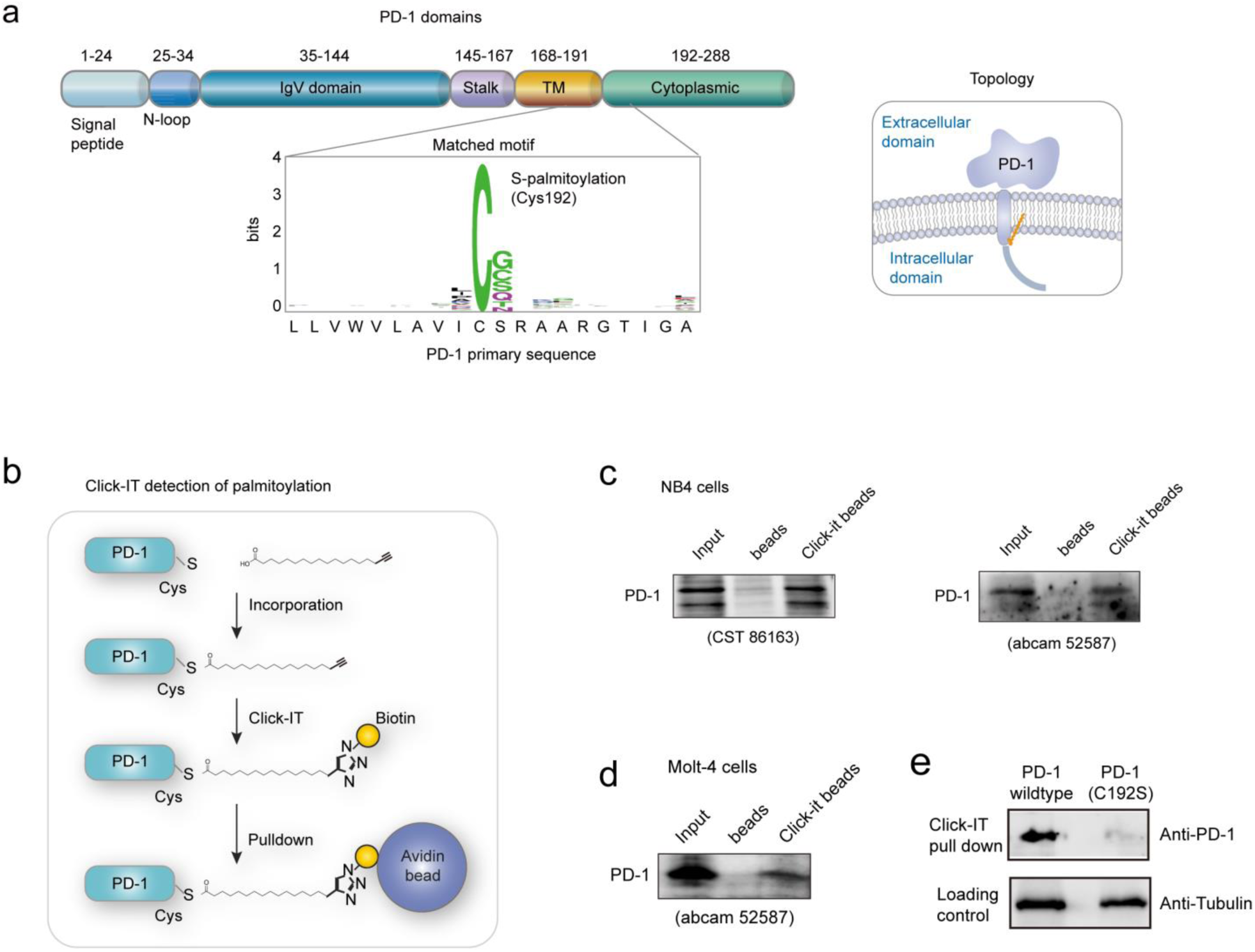
PD-1 is palmitoylated at C192. **a**. Prediction of PD-1 palmitoylation site at Cys192 by MDDPalm algorithm, with the matched motif in the insert and topology model on the right. **b.** Schematic representation of the Click-IT procedure used to detect PD-1 palmitoylated proteins. **c-d**. The Click-IT assay in NB4 (**c**) and Molt-4 (**d**) cells showing the same endogenous PD-1 band size in Click-IT beads compared to input lysate. The click-IT assay was performed on NB4 cells using an independent PD-1 antibody (right panel). The antibodies used for the final detection are indicated below the gels. **e**. The Click-IT was performed on cell transiently expressing wildtype PD-1 or Cys192Ser (C192S) mutated PD-1 to demonstrate that no palmitoylation was detected in the C192S PD-1 mutant.

### Palmitoylation of PD-1 stabilize its protein level

In general, palmitoylated protein “stabilize” protein level, suggesting that the primary function of palmitoylated PD-1 is to maintain a high protein level. As expected, PD-1 is abundantly expressed in many cancer cell lines (**Fig.2a**). To evaluate the impact of palmitoylation on the protein level of PD-1, we took advantage of 2-bromopalmitate (2-BP), a general inhibitor of palmitoylation ^15^. Treating A375 or PC3 cells significantly decrease PD-1 protein level as judged by immunofluorescence (**Fig. 2b**). Conversely, HepG2, LoVo and SW116 cells treated with Palmostatin-B (PalmB), a palmitoylation activator ^16^ significantly increase PD-1 protein expression (**Fig. 2c**). Thus, palmitoylation regulates the protein level of PD-1 and is one of the mechanisms by which cancer cells maintain a high level of PD-1.

**Figure 2.**
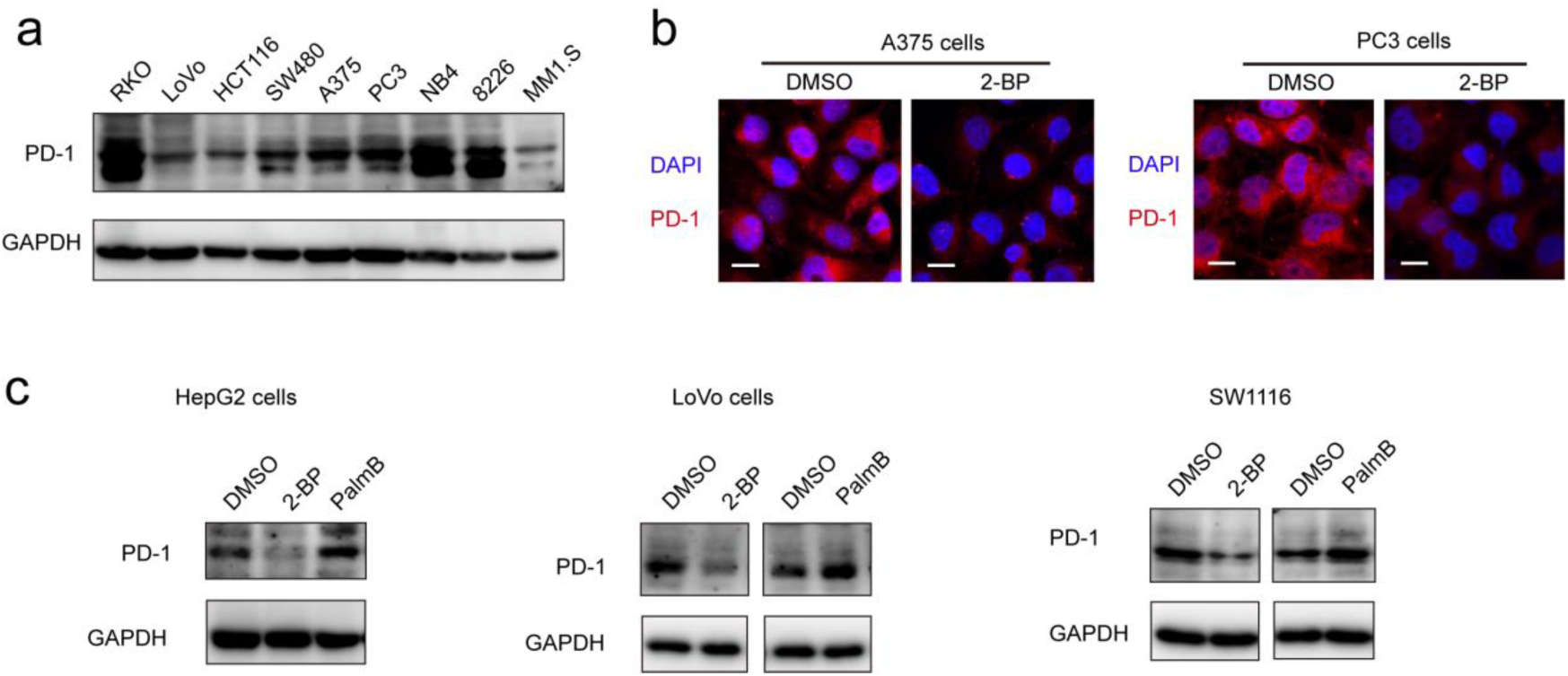
Palmitoylation of PD-1 stabilize its protein level. **a**. Expression of PD-1 in a panel of cancer cell lines by western-blot using anti-PD-1 specific antibody. **b**. Expression of PD-1 is reduced in cells treated with 2-BP (palmitoylation inhibitor) as shown by immunofluorescence. **c**. The treatment with 2-BP and PalmB (palmitoylation activator) respectively decreased and increased PD-1 expression.

### PD-1 is palmitoylated by DHHC9

Palmitoylation is an enzymatic reaction performed by the family of Asp-His-His-Cys (DHHC) acyltransferases ^17^. In order to identify the predominant DHHC family member responsible for PD-1 palmitoylation, we performed a systematic siRNA screen of the main DHHC family member in A375 cells (**Fig.3a**). Individual knockdown by two independent siRNA sequences effectively reduce expression of the targeted DHHC gene by at least 2-fold but only siRNAs against DHHC9 significantly decreased the expression of PD-1 (**Fig.3a**). Since PD-1 palmitoylation maintain PD-1 protein level, reducing the key factor palmitoylating PD-1 decrease PD-1 palmitoylation and hence, decrease its protein level. Similarly, knocking down DHHC9 in RKO and HCT-116 cell lines decrease PD-1 protein level (**Fig.3b**). Consistently, overexpression of DHHC9 increased PD-1 expression in MDA-MB-231, RKO and A375 cancer cell lines (**Fig.3c**). Interestingly, DHHC9 is one of the most expressed DHHC in tumor samples expressing high level of PD-1 (**Supplementary Fig. 1**). Altogether, these results pinpoint to DHHC9 as the major enzyme for PD-1 palmitoylation in cancer cells.

**Figure 3.**
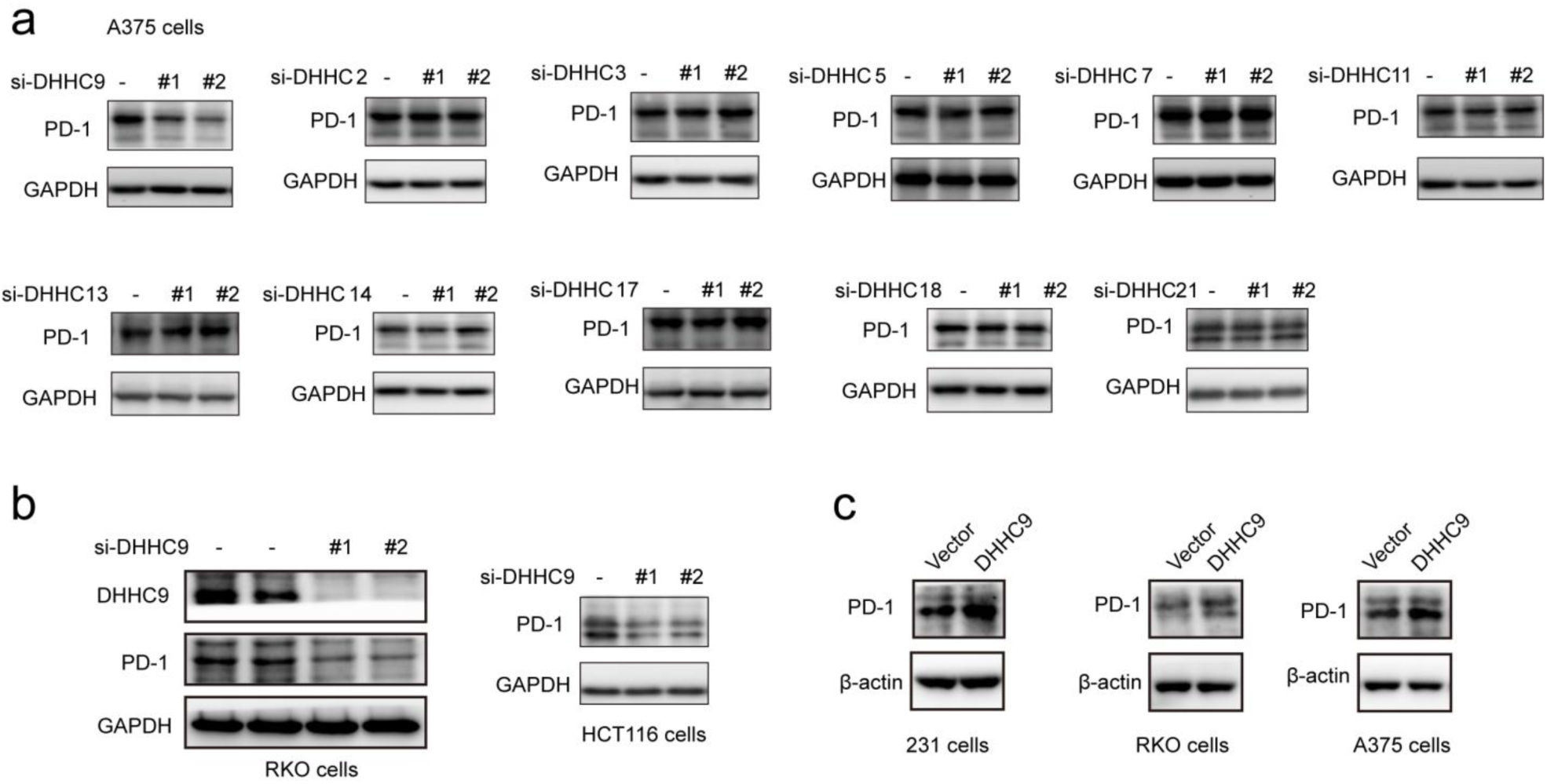
PD-1 is palmitoylated byDHHC9. **a.** Western blot showing the expression of PD-1 in A375 cells treated by two independent siRNAs targeting the indicated DHHC enzymes. **b**. Western blot showing the expression of PD-1 in RKO and HCT116 cells treated by two independent siRNAs targeting DHHC9. **c**. Immunoblot showing the expression of PD-1 in different cells overexpressing DHHC9.

### Palmitoylation promotes PD-1 binding to Rab11

Previous studies suggested that palmitoylation may affect protein stability by modulating its interaction with Rab11 and thereby its storage in recycling endosomes ^18^. Therefore, we performed co-IP assay by reciprocal pulldown of overexpressed PD-1 or its non-palmitoylated version (C192S) using anti-PD-1 or anti-RAB11 in HCT-1116, a colon cancer cell line expressing low PD-1 basal level. The results revealed that wild-type PD-1 but not the C192S mutant interacted with RAB11 (**Fig.4a&b**), supporting the importance of palmitoylation for PD-1:RAB11 interaction. Moreover, blockade of PD-1 palmitoylation by 2-BP also disrupted the PD-1:RAB11 interaction (**Fig.4c**). To confirm that interfering with the palmitoylation dependent PD-1:RAB11 interaction decreased the storage of PD-1 to recycling endosomes, we performed co-localization studies by immunofluorescence in two cell lines. As expected, PD-1 and RAB11 co-localized to recycling endosome in basal condition and upon treatment with 2-BP, the PD-1 signal intensity is reduced (**Fig.4d&e**). We further quantify this signal reduction by plotting the signal intensity of PD-1 and RAB11 over a representative endosome area (**Fig.4d&e, right panels**). Blocking palmitoylation by 2-BP also destabilized PD-1, which could be rescued by the lysosomal inhibitor chloroquine (**Fig.4f**). These findings consistently revealed that palmitoylation promotes the interaction between PD-1 and RAB11, thereby facilitating the transportation to recycling endosome and attenuating the degradation in the lysosome.

**Figure 4.**
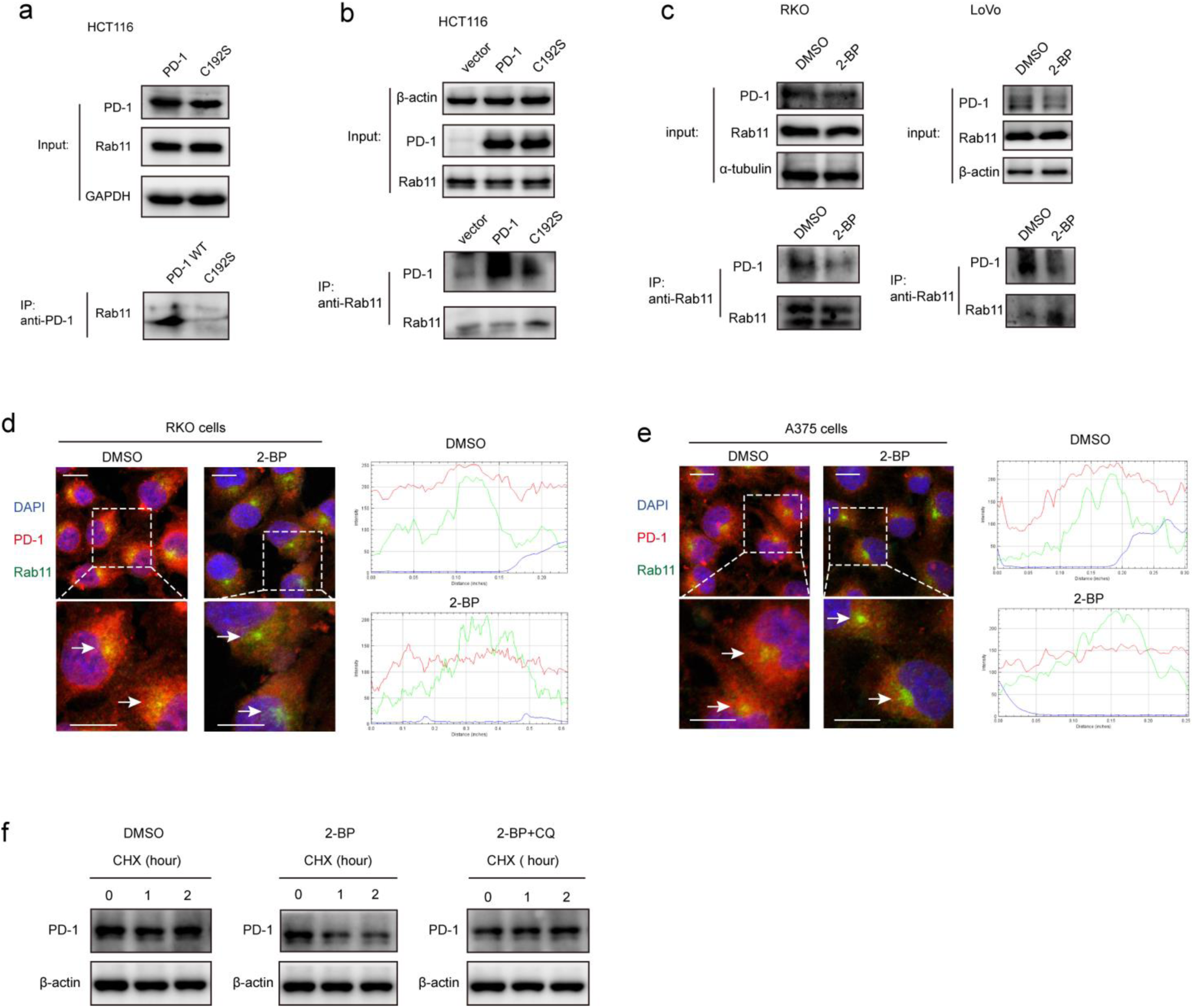
Palmitoylation promotes the binding of PD-1 to RAB11. **a**,**b.** The interaction between RAB11: wt PD-1 and RAB11: mutant PD-1 (C192S) are shown by co-IP in HCT-116 cells. The antibody for (**a**) PD-1 and (**b**) RAB11 was used for pulldown. **c.** The interaction between RAB11: wt PD-1 and RAB11: mutant PD-1 (C192S) are shown by co-IP in ROKO and LoVo cells. Disruption of palmitoylation by overexpression of PD-1 C192S or by 2-BP treatment significantly decreased the binding between PD-1 and RAB11, as shown by co-IP. **d**,**e**. Colocalization between PD-1 and RAB11 in (**d**) RKO and (**e**) A375 cells by immunofluorescence (right panels). White arrows indicate the regions of recycling endosome marked by RAB11, and the intensity profiles of PD-1 and RAB11 in the insert is quantified and plotted on the right. Upon 2-BP treatment, the enrichment of PD-1 on recycling endosome was abolished. **f**. The degradation of PD-1 was evaluated by cycloheximide (CHX)-chase assay. Treatment with 2-BP reduce PD-1 level and this decrease is rescued by the lysosomal inhibitor chloroquine (CQ).

### Palmitoylation promotes PD-1 cell autonomous mTOR signaling

Functionally, PD-1 expressing cancer cells possess growth and survival advantage that is independent of the adaptive immunity paracrine signaling with T cells ^7^. Mechanistically, it has been demonstrated that tumor intrinsic PD-1 functions through interactions with S6 and eIF4E, which are effectors of mTOR signaling ^5,6^. The binding to PD-1 increase the phosphorylation of S6 and eIF4E ^6^, leading to increased protein synthesis and cell proliferation as revealed by CCK8 assay. Thus, we hypothesize that PD-1 palmitoylation positively affect the binding of PD-1 to S6 and eIF4E. As predicted, there is an interaction between PD-1 and S6 as judged by co-IP (**Fig.5a**) in RKO cell line. Importantly, the suppression of PD-1 palmitoylation by DHHC9 depletion (**Fig.5b**), pharmacological inhibition using 2-BP (**Fig.5c**), or mutation of the Cys192 modification site (**Fig.5d**) effectively abolished the PD-1:S6 and PD-1:EIF4E interactions in LoVo, RKO and HCT-116 cell lines.

**Figure 5.**
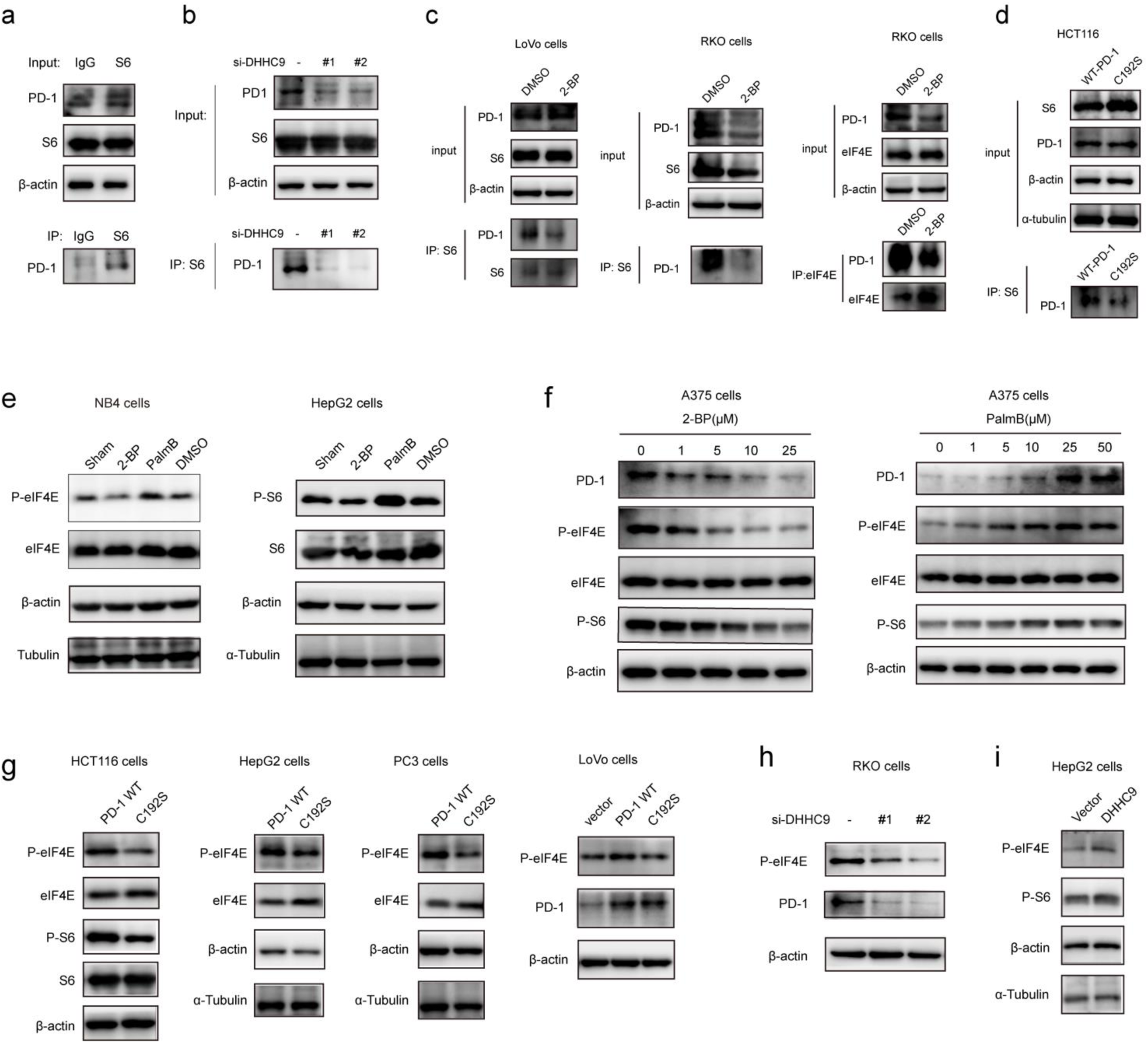
Palmitoylation promotes tumor-intrinsic PD-1 signaling. **a.** Interaction between endogenous PD-1 and S6 in tumor cells by co-IP. **b-d.** The interaction between PD-1 and S6/eIF4E were significantly decreased by depleting DHHC9 (**b**), 2-BP treatment (**c**) and mutating Cys192 (**d**) as shown by co-IP. **e** Modulating the expression level of PD-1 by 2-BP and PalmB impact on the phosphorylation level of the mTOR effectors eIF4E (left panels) and S6 (right panels). **f** Modulating the expression level of PD-1 by 2-BP and PB in a dose-dependent manner correlates with the phosphorylation of the mTOR effectors eIF4E (left panel) and S6 (right panel). **g** Overexpression of C192S decrease the phosphorylation of eIF4E and S6 in cancer cells. **h.** The phosphorylation of eIF4E and expression of PD-1 in RKO cells treated with siRNAs specific for DHHC9. **i.** The phosphorylation of eIF4E and S6 is increased in HepG2 cells overexpressing DHHC9.

Further, blocking palmitoylation by 2-BP decreased the phosphorylation of eIF4E (p-eIF4E) (**Fig. 5e**, left panel) while potentiating palmitoylation by Palmostatin-B increased p-eIF4E in NB4 cells. Similar results were obtained for S6 (p-S6) in HepG2 cells (**Fig. 5e**, right panel). The effects of modulating palmitoylation on the phosphorylation of S6 and eIF4E displayed dose-dependent effects (**Fig.5f**). Consistently, disrupting palmitoylation by mutating Cys192 residue also decreased the phosphorylation of eIF4E and S6 in HCT-116, HepG2, PC3 and LoVo cancer cell lines (**Fig.5g**) supporting our hypothesis that PD-1 palmitoylation regulates cancer cell autonomous mTOR signaling. Finally, siRNA-mediated depletion (**Fig.5h)** or upregulation (**Fig.5i**) of DHHC9 decreased and increased the phosphorylation of mTOR effectors, respectively. Thus, the palmitoylation dependent and cancer intrinsic function of PD-1 is to at least to signal through the mTOR pathway.

### Interfering with PD-1 palmitoylation reduce cancer cells growth in 2D and 3D in vitro

To evaluate cancer cells dependency on the palmitoylation status of PD-1 for cell growth, multiple cancer cell line expressing PD-1 were DHCC9 depleted then analyzed for growth rate by CCK8 assay. Compared to control condition, DHHC9 depletion significantly attenuated cancer cell growth (**Fig.6a**). Pharmacological blockade of palmitoylation by 2-BP decreased tumor growth (**Fig.6b**), while overexpressing wt PD-1 (PD-1 OE) but not C192S mutant (C192S OE) potentiated cancer cell growth (**Fig.6c&d**). Consistently, targeted depletion of DHHC9 expression also decreased the anchorage-independent proliferation (**Fig.6d**). Thus, interfering with PD-1 palmitoylation reduce cancer cell growth in vitro in 2D and 3D.

**Figure 6.**
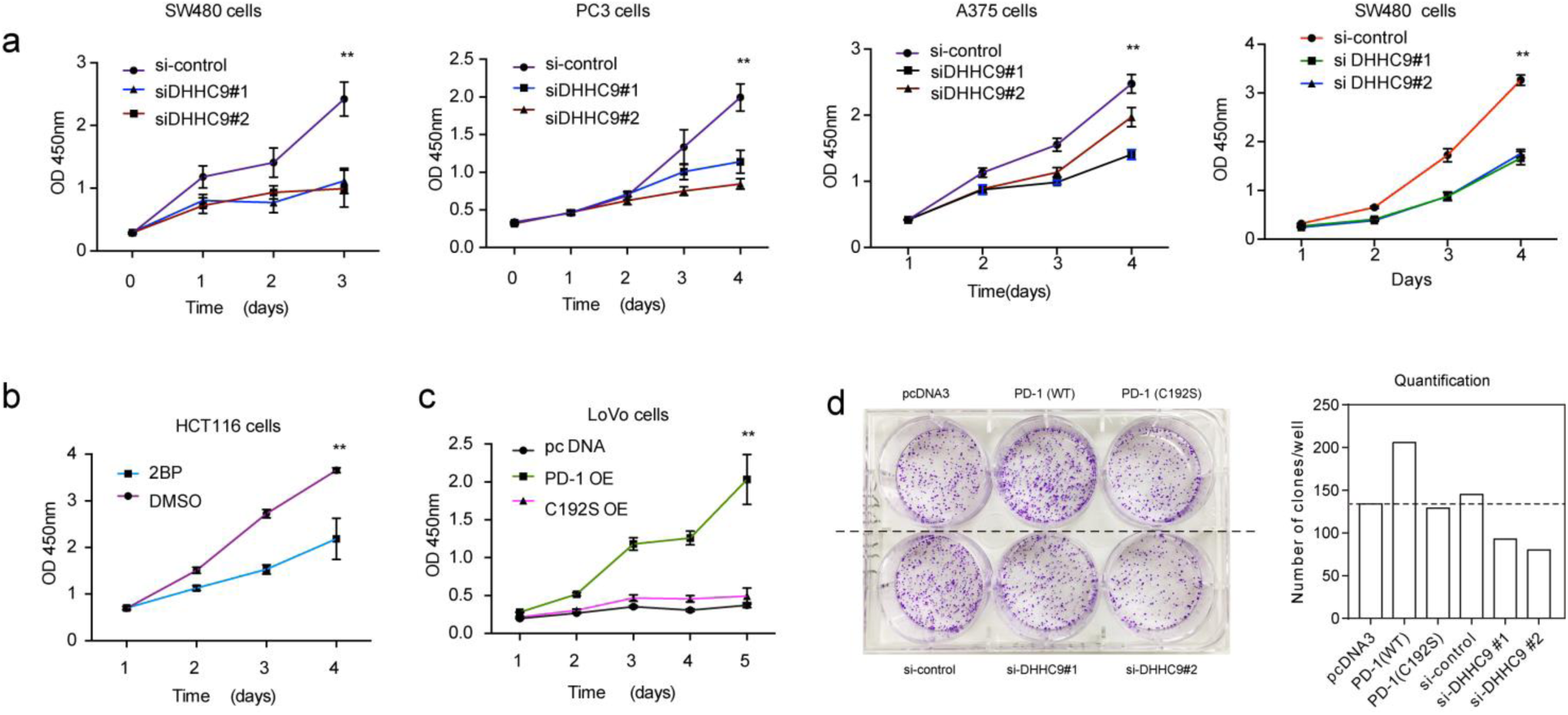
Targeting palmitoylation suppressed tumor-intrinsic PD-1 functions. **a**. CCK8 proliferation assay showing that the proliferation of cancer cells treated with siRNAs for DHHC9 is decreased. ** P<0.01, ANOVA test. **b.** Treatment of HCT-116 cells with 2-BP decrease the proliferation of cells as detected by CCK8 assay. **c.** Overexpressing the C192S mutant (C192S OE) decrease cell proliferation compared to wt PD-1 (PD-1 OE) in LoVo cells. **d.** Overexpressing wt PD-1 (PD-1 OE) increase anchorage-free colony formation assay and targeting DHHC9 decrease it compared to control conditions.

### A designed peptide targeting PD-1 palmitoylation and expression

Our group is successful at rationally designing peptide with biological activity and potential therapeutic benefit ^19^. To impair with PD-1 palmitoylation, we decided to design a peptide that interfere with the main PD-1 palmitoylation enzyme DHCC9 through competitive inhibition. To this end, we start by the identified sequence surrounding the palmitoylation site of PD-1 at Cys192 (**Fig.1a**) and subsequently optimized its cell-penetrating potential *in silico* using to CellPPD predictor ^20^. This led to the candidate peptide VICSRAAR (herein named as PD1-PALM), which shown in **Fig.7a**. To demonstrate its direct role in interfering with PD-1 palmitoylation, the Click-IT assay was used in cells treated with PD1-PALM. As expected, a lower level of PD-1 was retrieved in cells treated with PD1-PALM compared to control (**Fig.7b**). Accordingly, PD1-PALM decreased PD-1 expression in a dose-dependent manner in both SW480 and RKO cells (**Fig.7c**). Thus, PD1-PALM decrease PD-1 expression by interfering with its palmitoylation and may be an effective therapeutic strategy to target PD-1 intrinsic function in cancer cells.

**Figure 7.**
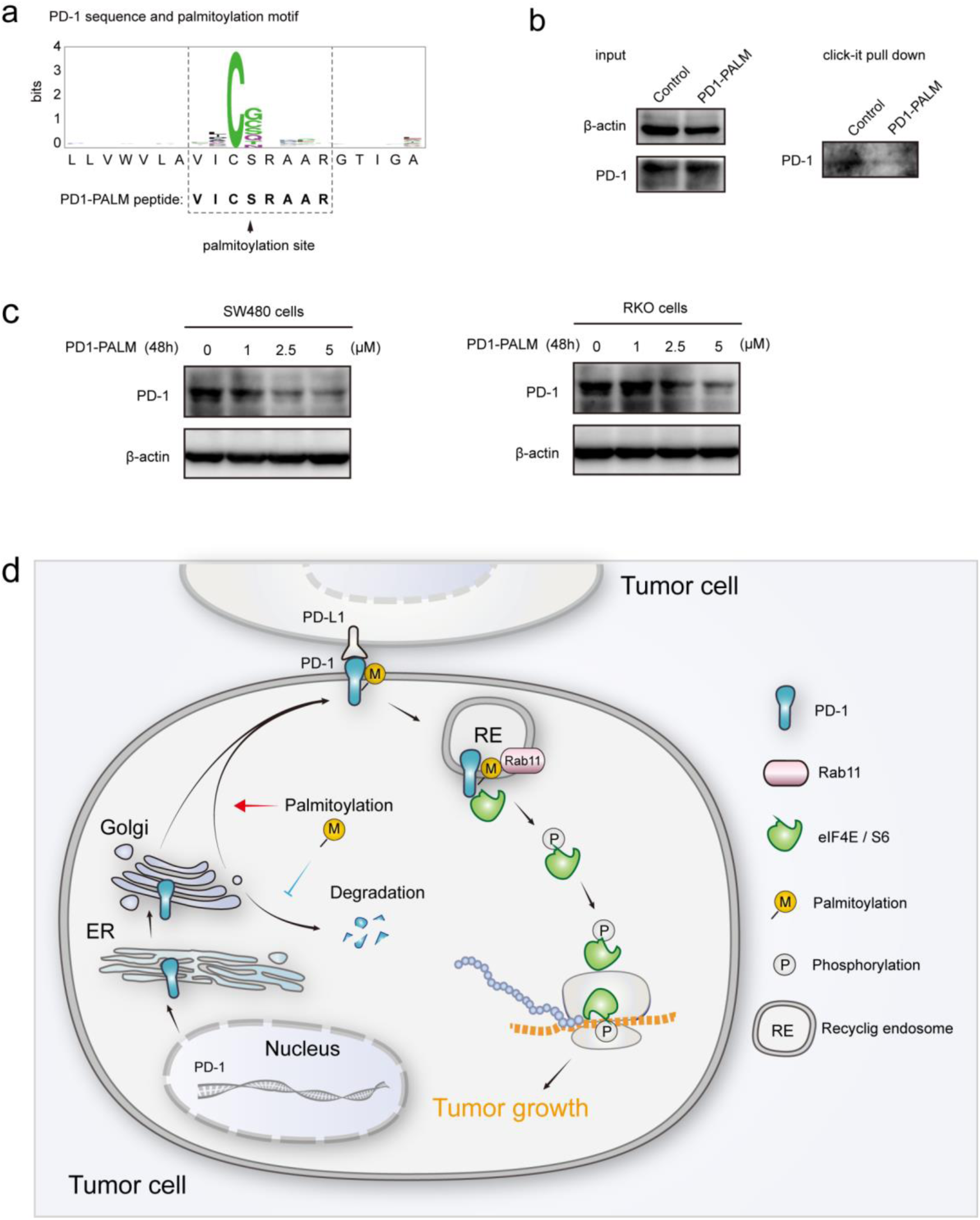
Competitive inhibitor of PD-1 palmitoylation. **a.** The sequence of the designed PD1-PALM peptide. **b.** Click-IT assay showing the decreased palmitoylation of PD-L1 in cells treated by the PD1-PALM peptide. The input is shown on the left, and PD-L1 detected in the Click-IT pulldown is shown on the right. **c.** The expression of PD-1 in SW480 (left panel) and RKO (right panel) cells treated by different concentrations of PD1-PALM peptide. **d.** Cancer intrinsic function and regulation of palmitoylated PD-1. Schematic model showing the roles of palmitoylation in promoting PD-1 expression and intrinsic mTOR signaling to facilitate tumor growth.

### Discussion

As a crucial target of tumor immunotherapy, the regulation of PD-1 in both immune cells and tumor cells is of prime importance in cancer immunology. For the first time, we report the palmitoylation of PD-1 and the role of this lipid modification in maintaining PD-1 expression and regulating its tumor intrinsic signaling.

Our results suggest that palmitoylation of PD-1 promotes its interaction with Rab11, which is a key molecule in transporting the cargo proteins to recycling endosome for storage. Blockade of palmitoylation decreased the trafficking of PD-1 to recycling endosome and promoted its lysosomal degradation (model in **Fig.7d**). More interestingly, palmitoylation of PD-1 also significantly enhanced the interaction between PD-1 and the effector proteins of mTOR signaling (S6 and eIF4E), thereby triggering the activation of mTOR signaling and facilitating tumor growth. Since the S6 and eIF4E proteins are cytosolic, it is not surprising that the palmitoylation of PD-1 on its cytosolic domain can regulate the interaction with the above mTOR effector proteins.

Prevalent antibody-based therapies can efficiently block PD-1 on cell surface ^21^, but PD-1 is also maintained in the recycling endosome and other subcellular compartments. The intracellular PD-1 may be redistributed to cell surface, and potentially compromise the effects of therapeutic antibodies. Therefore, it is a potentially more persistent strategy to deplete PD-1 in the whole cell. In the present study we found that blocking palmitoylation can efficiently deplete the expression of PD-1, thereby presenting an alternative strategy for targeting PD-1. In addition to addressing the importance of palmitoylation on tumor-intrinsic PD-1, our study also identified the main enzyme responsible for PD-1 palmitoylation in cancer cells. Given the importance of PD-1 in both cancer cells and immune cells, simultaneously suppressing PD-1 in both tumor cells and immune cells may be an effective strategy.

In summary, our results highlight an important role of palmitoylation in regulating PD-1, and provide additional opportunities for targeting PD-1 as a crucial target for tumor immunotherapy.

## Methods

### Cell culture and transfection

The human colorectal cancer HCT-116, LOVO, RKO, SW480, A375, PC3, NB4, 8226 and MM1.S cells were purchased from ATCC. All cell lines were tested and verified to be free of Mycoplasma. Cells were maintained in RPMI 1640 or DMEM or Myco 5A medium (Gibco, Gaithersburg, MD, USA) supplemented with 10% (vol/vol) fetal bovine serum (Invitrogen, Carlsbad, CA, USA) and cultured in a humidified incubator at 37°C under 5% CO2. For transfection, cells were seeded at 50% confluence 24 h before transfection, and then transfected using FuGENE HD (Promega, Fitchburg, WI, USA) according to the product manual. Briefly, the transfection complex was made by 1 μg plasmid, 3 μl FuGENE HD and 100 μl OptiMEM. For transfection of siRNAs, the sequences of siRNAs are shown in **Supplementary Table S1**. Six hours after the complex was added to the cells, normal culture media was used to culture cells for additional 48 h.

### Antibodies and chemicals

The primary antibodies for HA tag (#3724, CST), RAB11 (610656, BD), PD-1 (#86163, CST and #52587, Abcam, AF1086,R&D,#192106,R&D) were commercially available. Secondary antibodies used in immunofluorescence assay were: AF488-anti-Mouse (Invitrogen), AF594-anti-rabbit (Invitrogen), AF488-anti-rabbit (Invitrogen), AF594-anti-mouse (Invitrogen). Small molecular compounds such as 2-BP (Sigma), PalmostatinB (Millipore), Cycloheximide (C1988, Sigma), Chloroquine (C6628, Sigma), Click-iTpalmiticacid, azide (C10265,Thermo Scientific), Click-iT Cell Reaction Buffer Kit (C10269,Thermo Scientific), Biotin-alkyne (Sigma),Streptavidin (Sepharose Bead Conjugate) (#3419,CST),Goat anti-Human IgG (H+L) Cross-Adsorbed Secondary Antibody, Alexa Fluor 488(A11013, Thermo Scientific) and DAPI (0100-20, Southern Biotech) were also purchased from the indicated suppliers.

### Plasmids construction

The expression vectors encoding pcDNA3.1-Flag-PD-1 and pcDNA3.1-DHHC9 were generated by inserting synthesized cDNAs into the pcDNA3.1 vector. PD-1 C192S mutant was generated by site-directed mutagenesis PCR reaction using platinum PWO SuperYield DNA polymerase (Roche, Basel, Switzerland) according to the product manual. All plasmids were sequenced to confirm if the designed mutation is present, without any other unwanted mutation.

### Click-iT identification of PD-L1 Palmitoylation

48 hours after transfection of PD-1 or PD-1 C192S mutant, 100μM of Click-iTpalmitic acid-azide was added to the cell medium with gentle mix, and then incubated at 37°C, 5% CO2 for 6 hours. After 6 hours incubation, the medium was removed and the cells was washed three times with PBS before the addition of lysis buffer (1% SDS in 50mM Tris-HCl, PH 8.0) containing protease and phosphatase inhibitors at appropriate concentrations. We incubated the cells for 20 min on ice, then tilt the plates and pipet the lysate into a 1.5 mL microcentrifuge tube. Then we sonicated the lysate with a probe sonicator to solubilize the proteins and disperse the DNA. After vortexing the lysate for 5 minutes and centrifuging the cell lysate at 13,000–18,000 × g at 4°C for 5 minutes, we transferred the supernatant to a clean tube and determine the protein concentration using the EZQ® Protein Quantitation Kit (Cat. no. R33200,Thermo Scientific) or another method. Thus, the protein sample was reacted with biotin-alkyne using the Click-iT Protein Reaction Buffer Kit (Cat. no. C10276, Thermo Scientific) following protocols in the instruction sheet. Then biotin-alkyne-azide-plamitic-protein complex was pulled down by Streptavidin and after washing, the pellets were subjected to immunoblotting detection for PD-L1.

### Immunofluorescence

Cells were fixed with 4% PFA for 20 min, permeated as well as blocked with PBS buffer containing 1%BSA and 0.2% Triton-100 for 1 hour at room temperature. The cells were incubated with the primary antibody at 4°C overnight. After washing 3 times with PBS, the cells were incubated with the fluorescent conjugated secondary antibody for 30min at room temperature. The slides were mounted with Prolong Gold and imaged on Zeiss 710 confocal microscope.

### Immunoblotting and immunoprecipitation

For immunoblotting, cells were lysed either in 1% Triton X-100 in TBS pH7.6 with Roche complete protease inhibitor for 30 min on ice followed by pelleting of insoluble material by centrifugation. Lysates were heated to 100 °C in SDS sample buffer with 50 mM DTT for 10 min, separated by SDS–PAGE, and transferred to PVDF membrane (Millipore). Membranes were blocked in 5% BSA in TBS, probed with the indicated antibodies, and reactive bands visualized using West Pico (Thermo Fisher Scientific).

For co-immunoprecipitation experiments, cells were lysed in IP buffer (#87787, Thermo Scientific) plus Roche complete protease inhibitor for 10 min on ice followed by the addition of benzonase (Sigma) for 25 min at room temperature. Then the lysate was centrifuged at 15000 rpm at 4°C to remove the precipitation. Then the supernatants were incubated with primary antibody at slow roating speed at 4 °C overnight, followed by addition of protein A or protein G agarose beads and incubation for a further 2 h at 4 °C. After four washes in PBST(PBS with 0.01% Tween 20), samples were eluted in SDS sample buffer with 50 mM DTT for 10 min at 100 °C, separated by SDS–PAGE and immunoblotted as described.

### Detection of PD-1 degradation rate

After the indicated treatments, cells were incubated with cycloheximide (50μg/mL) for different time-point. And then the cells were subjected to immunoblotting assay and the immunoblotting images were quantified by Quantity One software.

### Cell Proliferation Assay

Cell proliferation assay was done using a Cell Counting Kit-8 (CCK-8) (Dojindo), following manufacturer’s instructions. Twenty-four hours after treatments, cells were harvested and seeded into 96-well plates at an initial density of 2,000 cells per well in 100 μl culture medium. At the indicated time points for analysis, 10 μL of CCK-8 solution was added to each well with serum-free medium, followed by incubation for 2 hours in a humidified incubator at 37°C with 5% CO2 supplement. The optical density (OD) at 450 nm (indicating formation of formazan) was measured using VERSA Max microplate reader (MDS Analytical Technologies) with SoftMaxPro software.

### Statistical analysis

Data in bar graphs indicate mean ± SD fold change in relation to control groups of 3 independent experiments. Statistical analyses were performed using SPSS (IBM). All the proliferation results containing more than two conditions were analyzed using ANOVA post hoc test (Tukey, compare all pairs of conditions). Statistical significance was considered when P value was below 0.05.

### Author contributions

HY, CL, JPB, HW, HL, CF, HS, JL and JX. contributed to the design and /or execution of experiments. HY, JPB and JX wrote the paper. JX conceived and supervised the study.

## Acknowledgements

This project was supported by grants from the National Key Research & Development (R&D) Plan (2016YFC0906000,2016YFC0906002); National Natural Science Foundation of China (81572326, 81322036, 81421001); Top-Notch Young Talents Program of China (ZTZ2015-48); “Tang Scholar” program (JX-2017); Shanghai Municipal Education Commission-Gaofeng Clinical Medicine Grant Support (20152514); “ShuGuang” project supported by Shanghai Municipal Education Commission and Shanghai Education Development Foundation (15SG16); National Key Technology Support Program (2015BAI13B07) to Jie Xu; The National Science Foundation of China (21335002) to Haojie Lu, and Natural Science Foundation of Shanghai (18ZR1402800) to Caiyun Fang. The funders had no role in study design, data collection and analysis, decision to publish, or preparation of the manuscript.

